# Gene co-expression networks in whole blood implicate multiple interrelated molecular pathways in obese asthma

**DOI:** 10.1101/181651

**Authors:** Damien C. Croteau-Chonka, Zhanghua Chen, Kathleen C. Barnes, Albino Barraza-Villarreal, Juan C. Celedón, W. James Gauderman, Frank D. Gilliland, Jerry A. Krishnan, Andrew H. Liu, Stephanie J. London, Fernando D. Martinez, Joshua Millstein, Edward T. Naureckas, Dan L. Nicolae, Steven R. White, Carole Ober, Scott T. Weiss, Benjamin A. Raby

**Affiliations:** Channing Division of Network Medicine, Department of Medicine, Brigham and Women’s Hospital and Harvard Medical School, Boston, MA, USA; Division of Environmental Health, Department of Preventive Medicine, Keck School of Medicine, University of Southern California, Los Angeles, CA, USA; Division of Biomedical Informatics and Personalized Medicine, Department of Medicine, University of Colorado School of Medicine, Anschutz Medical Campus, Aurora, CO, USA; National Institute of Public Health, Cuernavaca, Morelos, México; Division of Pulmonary Medicine, Allergy and Immunology, Children’s Hospital of Pittsburgh of the University of Pittsburgh Medical Center, University of Pittsburgh, Pittsburgh, PA, USA; Division of Biostatistics, Department of Preventive Medicine, Keck School of Medicine, University of Southern California, Los Angeles, CA, USA; Division of Pulmonary, Critical Care, Sleep, and Allergy, Department of Medicine, University of Illinois at Chicago, Chicago, IL, USA; Division of Allergy and Clinical Immunology, Department of Pediatrics, National Jewish Health and University of Colorado School of Medicine, Denver, CO, USA; Division of Intramural Research, Department of Health and Human Services, National Institute of Environmental Health Sciences, National Institutes of Health, Research Triangle Park, NC, USA; Arizona Respiratory Center and BIO5 Institute, University of Arizona, Tucson, AZ, USA; Section of Pulmonary and Critical Care Medicine, Department of Medicine, University of Chicago, Chicago, IL, USA; Department of Human Genetics, University of Chicago, Chicago, IL, USA; Section of Genetic Medicine, Department of Medicine, University of Chicago, Chicago, IL, USA; Department of Statistics, University of Chicago, Chicago, IL, USA; Partners HealthCare Personalized Medicine, Partners Health Care, Boston, MA, USA; BWH Pulmonary Genetics Center, Division of Pulmonary and Critical Care Medicine, Department of Medicine, Brigham and Women’s Hospital and Harvard Medical School, Boston, MA, USA

**Author notes:** Corresponding Author Damien C. Croteau-Chonka, Ph.D. 181 Longwood Avenue Boston, MA 02115-5804.

**Keywords:** asthma, obesity, gene expression, blood, inflammation, platelet, integrin, smooth muscle, extracellular matrix, Hedgehog signaling

## Abstract

**Background:** Asthmatic children who develop obesity have poorer outcomes compared to those that do not, including poorer control, more severe symptoms, and greater resistance to standard treatment. Gene expression networks are powerful statistical tools for characterizing the underpinnings of human disease that leverage the putative co-regulatory relationships of genes to infer biological pathways altered in disease states.

**Objective:** The aim of this study was to characterize the biology of childhood asthma complicated by adult obesity.

**Methods:** We performed weighted gene co-expression network analysis (WGCNA) of gene expression data in whole blood from 514 adult subjects from the Childhood Asthma Management Program (CAMP). We then performed module preservation and association replication analyses in 418 subjects from two independent asthma cohorts (one pediatric and one adult).

**Results:** We identified a multivariate model in which four gene co-expression network modules were associated with incident obesity in CAMP (each *P* < 0.05). The module memberships were enriched for genes in pathways related to platelets, integrins, extracellular matrix, smooth muscle, NF-κB signaling, and Hedgehog signaling. The network structures of each of the four obese asthma modules were significantly preserved in both replication cohorts (permutation *P* = 9.999E-05). The corresponding module gene sets were significantly enriched for differential expression in obese subjects in both replication cohorts (each *P* < 0.05).

**Conclusions:** Our gene co-expression network profiles thus implicate multiple interrelated pathways in the biology of an important endotype of obese asthma.

**Key Messages:**

- We hypothesized that individuals with asthma complicated by obesity had distinct blood gene expression signatures.
- Gene co-expression network analysis implicated several inflammatory biological pathways in one form of obese asthma.

**Capsule Summary:** This work addresses a knowledge gap about the molecular relationship between asthma and obesity, suggesting that an endotype of obese asthma, known as asthma complicated by obesity, is underpinned by coherent biological mechanisms.

**Abbreviations:**

CAMP: Childhood Asthma Management Program
WGCNA: weighted gene co-expression network analysis
Asthma BRIDGE: Asthma BioRepository for Integrative Genomic Exploration
GACRS: Genetics of Asthma in Costa Rica Study
CHS: Southern California Children’s Health Study
BMI: body mass index
BIC: Bayes Information Criterion
HUGO: Human Genome Organisation
PC: principal component
GSEA: gene set enrichment analysis
IL-1: interleukin-1
Hh signaling: Hedgehog signaling

## Introduction

Obesity is recognized as an important risk factor for, and a major co-morbidity of, asthma in both children and adults [1, 2]. Obesity is associated with a significantly higher risk of developing asthma, worse asthma symptoms, poor asthma control, and greater resistance to standard asthma therapies [1]. A recent review [2] postulates at least three (likely four) endotypes of obese asthma [3, 4], distinguished by the relative timings of each disease’s onset (onset during childhood versus adulthood) and by their underlying pathophysiologies (differing effects of obesity on airway structure, function, sensitivity, and/or inflammation). A more complete understanding of the complex molecular relationships between obesity and asthma would drive the development of specific and effective preventative and therapeutic strategies for this overlap syndrome.

Despite large numbers of published studies characterizing global gene expression of each condition separately, there are few examples of comprehensive blood transcriptional profiling studies assessing the interrelationships of asthma and obesity. Of the three published studies [5-7] to date, only one [5] directly compared gene expression profiles between obese and non-obese asthmatic individuals; it was of small sample size (*n* < 10 per group) and lacked replication. Furthermore, all three studies were cross-sectional, making it difficult to pinpoint which endotypes of obese asthma they were characterizing. With these limitations, the three studies cumulatively implicated pathways related to the adaptive and innate immune systems in obese asthma. Larger, longitudinal studies are needed to determine the generalizability and reproducibility of these findings.

To address these knowledge gaps, we performed weighted gene co-expression network analysis (WGCNA) [8] in a cohort of asthmatic children who were followed longitudinally into young adulthood. Gene co-expression networks are powerful statistical tools for characterizing the underpinnings of human disease that use the putative co-regulatory relationships of genes to infer the biological pathways altered in disease states. We studied children who participated in the Childhood Asthma Management Program (CAMP), a four-year clinical trial and 12-year natural history follow-up study of the effects of inhaled anti-inflammatory medications in patients with mild to moderate persistent asthma [9]. A prior analysis in CAMP showed that asthmatic children who became obese during the observation period had impaired growth in lung function compared to those who did not [10]. To determine whether these children had a distinct molecular profile that could contribute to this association, we performed WGCNA using gene expression data from peripheral blood collected at early adulthood available from a subset of 514 CAMP participants, including 104 (20%) that became obese by adulthood (incident obesity) and then replicated in two additional, independent asthma cohorts (one pediatric and one adult, totaling 418 subjects).

Through this effort, we discovered a set of four gene co-expression network modules that were reproducibly associated with obesity in asthma. These modules were enriched for genes linked to several important molecular processes, implicating platelet, integrin, extracellular matrix, smooth muscle, NF-κB signaling, and Hedgehog (Hh) signaling pathways in the biology of obese asthma.

## Materials and Methods

### Asthma study cohorts

The CAMP study was initiated as a randomized clinical trial studying the long-term effects of three asthma medications on disease progression [9] (clinicaltrials.gov identifier NCT00000575). Subjects in this cohort had mild-to-moderate asthma and were phenotyped longitudinally from childhood (aged five to 12 years at baseline of the trial) until early adulthood, as previously described [10].

Replication was performed in two independently ascertained asthma cohorts: The Asthma BioRepository for Integrative Genomic Exploration (Asthma BRIDGE) and the Genetics of Asthma in Costa Rica Study (GACRS). Asthma BRIDGE consists of an ethnically diverse set of asthmatic adults from the United States who were studied as part of the EVE asthma genetics consortium [11] and who were further characterized with blood gene expression profiles [12]. For the current study, we used a subset of adult Asthma BRIDGE subjects from the Southern California Children’s Health Study (CHS) for whom both longitudinal asthma and obesity phenotypes were available, spanning childhood (aged 5 to 8 years of age at study enrollment) until early adulthood [13]. GACRS consists of asthmatic children (aged six to 14 years) ascertained from the Central Valley of Costa Rica, a population isolate with a high prevalence of asthma [14]. Subjects in GACRS had only cross-sectional data on asthma and obesity phenotypes.

This analysis of human subjects data was approved by the Institutional Review Board of Brigham and Women’s Hospital.

### Gene expression data

The cross-sectional CAMP gene expression data have been previously described [12]. Briefly, 620 CAMP subjects with whole blood samples collected only at early adulthood were assayed with the Illumina HT-12 (version 3) platform with 47,009 high-quality probes. These expression data were quantile-normalized and log_2_-transformed. A set of 106 subjects who were obese at enrollment in CAMP (body mass index (BMI) ≥ 95^th^ percentile for age and sex) were excluded to focus the analysis on asthma complicated by incident obesity [10]. Following the exclusion of probes with low variances [15], the final dataset comprised 514 subjects and 10,448 probes. For replication, cross-sectional whole blood gene expression data were also assayed with the Illumina HT-12 array (version 4): 47,009 high-quality probes for 91 adult subjects in CHS [12]; and 47,256 high-quality probes for 329 pediatric subjects in GACRS (Virkud *et al.*, unpublished). A subset of 10,448 probes in CHS and 10,437 probes in GACRS were matched to the CAMP data by their universal nucleotide identifiers [16].

### Statistical analyses

Analyses were performed with Bioconductor in R (both version 3.3). We performed WGCNA on each gene expression dataset with the R package “WGCNA” [8] (version 1.51). The fundamental premise of WGCNA is that a scaled matrix of all pair-wise correlations among measured genes can be used to model how those genes act in coherent regulatory communities that map to shared biological functions. In mathematical terms, certain groups of genes are more strongly correlated with each other than with all other genes based on absolute (unsigned) values of pair-wise correlations of expression. We first characterized unsigned correlation networks and their relationships with each other [17] in the discovery cohort (CAMP) and later examined the properties of those same networks in the replication cohorts (CHS and GACRS).

We next tested the association of all network modules with adult obesity case status using a multivariate linear model. We used the “glmulti” package (version 1.0) to perform automated model selection based on a genetic algorithm [18]. Rather than exhaustively examining all possible combinations of model predictors, which can be a very large number, this algorithm randomly explores the model search space while biasing its search towards more informative models. The algorithm generates an initial random population of possible models, evaluates the fitness of each one, and then recombines and mutates those models to join more informative predictors to create the next generation of models. This feature-mixing process repeats until the mean and highest fitness of the model population do not substantially change between steps. Models were ranked based on Bayes Information Criterion (BIC). The parsimonious association model was further adjusted for possible demographic and technical confounders: age, sex, self-reported race/ethnicity, and measured blood cell counts (% of eosinophils, monocytes, neutrophils, and lymphocytes). Blood cell counts were imputed for CHS and GACRS based on gene expression using the R package “CellMix” (version 1.6) [19]. Eosinophil counts could not be imputed with the default CellMix reference panel, but were directly measured in both replication cohorts.

To assign biological interpretability to the associated modules from the discovery cohort, we performed pathway overrepresentation analyses using g:Profiler [20] (database version: r1730_e88_eg35) and the corresponding R package “gProfiler” (version 0.6). Human Genome Organisation (HUGO) gene symbols served as gene identifiers for each expression probe. To increase the specificity in the enrichment results, we set a more stringent threshold of overlap (intersection of query and test sets ≥ 3) and excluded consideration of inferred electronic annotations (i.e., annotations created by *in silico* curation methods of potentially lower quality). An adjusted *P* value was calculated in a manner that accounted for the hierarchical relationships among the tested gene sets [20].

Replication analysis in CHS and GACRS consisted of two steps: module preservation and replication of differential gene expression. We tested network module preservation between the discovery and replication cohorts using NetRep (version 1.0) [21]. For each replication cohort, 10,000 network permutations were used to determine preservation significance. Replication differential expression analysis and gene set enrichment analysis (GSEA) [22] was performed using the R/Bioconductor package “phenoTest” (version 1.18). This two-step replication analysis served to complement the module association approach used for the discovery cohort. The differential expression model included the same demographic and technical covariates as the network association model tested in the discovery cohort. Gene set enrichment was assessed using a Wilcoxon test [22], which addressed whether the average value of the genes that belonged to a given set was different from the average value of the remaining genes that did not belong to that set. We chose the memberships of the four co-expression modules as specific gene sets we hypothesized to be differentially expressed in obese subjects.

## Results

### Gene co-expression network modules of obese asthma in discovery cohort

The aim of our study was to better understand the biological processes underlying the relationship of obesity with asthma. For this, we used gene expression profiling of the peripheral blood of adolescents with asthma, comparing those who did (*n* = 104) and did not (*n* = 410) become obese over approximately 16 years of observation (**Figure 1**). Obesity during adulthood was defined as a body mass index (BMI) ≥ 30. All subjects were non-obese during childhood (BMI < 95^th^ percentile).

**Figure 1:**
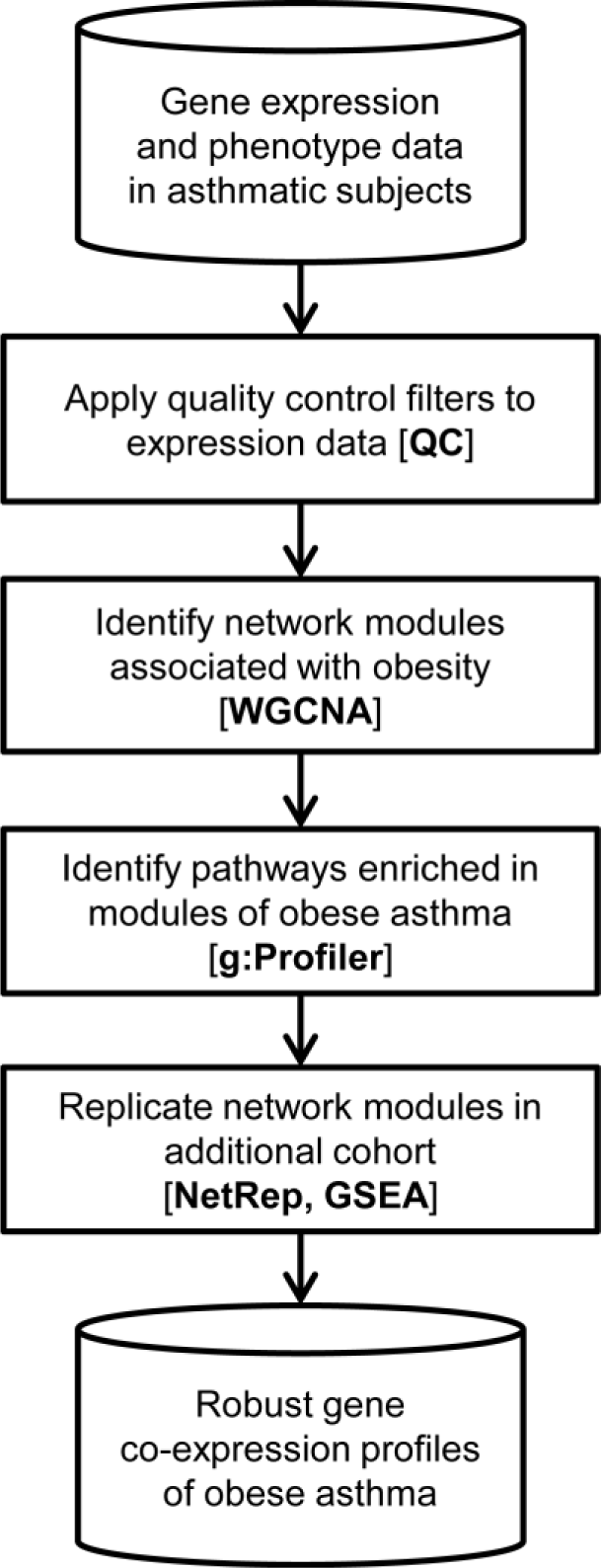
Analysis overview. In this schematic of this gene expression study of obese asthma, cylinders represent inputs and outputs, and boxes represent analysis steps. More detail for each analysis stage can be found in the Methods section. QC, quality control; WGCNA, weighted gene co-expression network analysis; GSEA, gene set enrichment analysis.

Baseline characteristics for the 514 subjects with both complete phenotype and expression data are provided in **Table 1**. Similar to patterns observed in the full CAMP cohort (including those without gene expression data) [10], we confirmed that, compared to children who did not develop obesity during the observation period, those who did become obese had higher BMIs at enrollment and included more subjects of non-European ethnicity (*P* < 0.05). The increased mean childhood BMI among CAMP subjects who became obese adults (78^th^ versus 53^rd^ percentile) suggested that they already bore an increased obesity susceptibility burden prior to the time point examined in this gene expression study. Though children who became obese were older at enrollment, we found no difference in age at the time of blood draw for gene expression (*P* > 0.1).

**Table 1:** CAMP discovery cohort characteristics. To test for differences between obese and non-obese subjects in the CAMP cohort, Welch two sample *t*-tests were used for continuous traits, and chi-squared tests were used for proportions.

|  | **Obese** | **Non-Obese** | **Difference *P*** |
| --- | --- | --- | --- |
| *N* | 104 | 410 | - |
| **Childhood BMI percentile** | 77.8 (15.5) | 53.1 (26.3) | 4.18E-28 |
| **Adult BMI** | 31.1 (3.9) | 23.5 (3.3) | 4.30E-39 |
| **Age at asthma onset (S.D.) [yrs.]** | 3.6 (2.6) | 3.9 (2.4) | 0.27 |
| **Age at study enrollment (S.D.) [yrs.]** | 9.1 (2.0) | 8.7 (2.1) | 0.09 |
| **Age at gene expression (S.D.) [yrs.]** | 21.2 (2.1) | 20.8 (2.2) | 0.15 |
| **% Female** | 37.5 | 38.0 | 1.00 |
| **Race/Ethnicity *N* (%)** |  |  |  |
| **European** | 67 (64.4) | 294 (71.7) | 0.02 |
| **Black/African American** | 18 (17.3) | 53 (12.9) |  |
| **Hispanic/Latino** | 16 (15.4) | 31 (7.6) |  |
| **Other** | 3 (2.9) | 32 (7.8) |  |
| **Log<sub>10</sub> IgE** | 2.5 (0.6) | 2.6 (0.6) | 0.28 |
| **% Eosinophils (S.D.)** | 3.8 (2.7) | 4.1 (2.9) | 0.29 |
| **% Lymphocytes (S.D.)** | 30.8 (7.5) | 31.6 (8.1) | 0.40 |
| **% Monocytes (S.D.)** | 7.5 (2.3) | 7.6 (2.2) | 0.54 |
| **% Neutrophils (S.D.)** | 57.5 (8.8) | 56.0 (9.4) | 0.18 |

Given the more adverse features of asthma observed in obese (vs. non-obese) CAMP subjects [1], we hypothesized that asthmatic subjects who became obese by adulthood would demonstrate coordinated patterns of global expression that were different from those observed in subjects who remained lean, and that the observed differences would implicate specific biological pathways that underlie differences between obese and non-obese asthma. To test this hypothesis, we first performed WGCNA on the adult whole blood expression data available in CAMP (**Figure 1**) [8]. To emphasize the impact of strong correlations over weak ones in the network construction, we chose an empirical soft threshold of 8, representing a strong model fit for scale-free topology (*R*^2^ > 0.8), which resulted in a mean connectivity of 10 genes (**Figure E1**).

Of the 10,448 genes considered in our analysis, 7,778 clustered into 17 distinct network modules of co-expressed genes (**Table E1**, **Table E2**, and **Figure E2**), which were each assigned an arbitrary color name. The remaining 2,670 genes that could not be assigned membership to a module because of insufficient co-expression were grouped as a collection termed the “null” module. Each module was represented by a summary eigengene (i.e., the first and largest principal component (PC) of variation among the co-expressed genes comprising that module). Correlation analysis of these eigengenes revealed evidence for higher-order structuring of the modules’ relationships to each other (**Figure 2**). Of the 17 eigengenes, nine were highly correlated with each other and formed a meta-module (based on average linkage hierarchical clustering) [8]; a second group of four eigengenes formed a smaller meta-module; and the four remaining modules exhibited sufficiently less correlation with any others and were not considered a distinct group.

**Figure 2:**
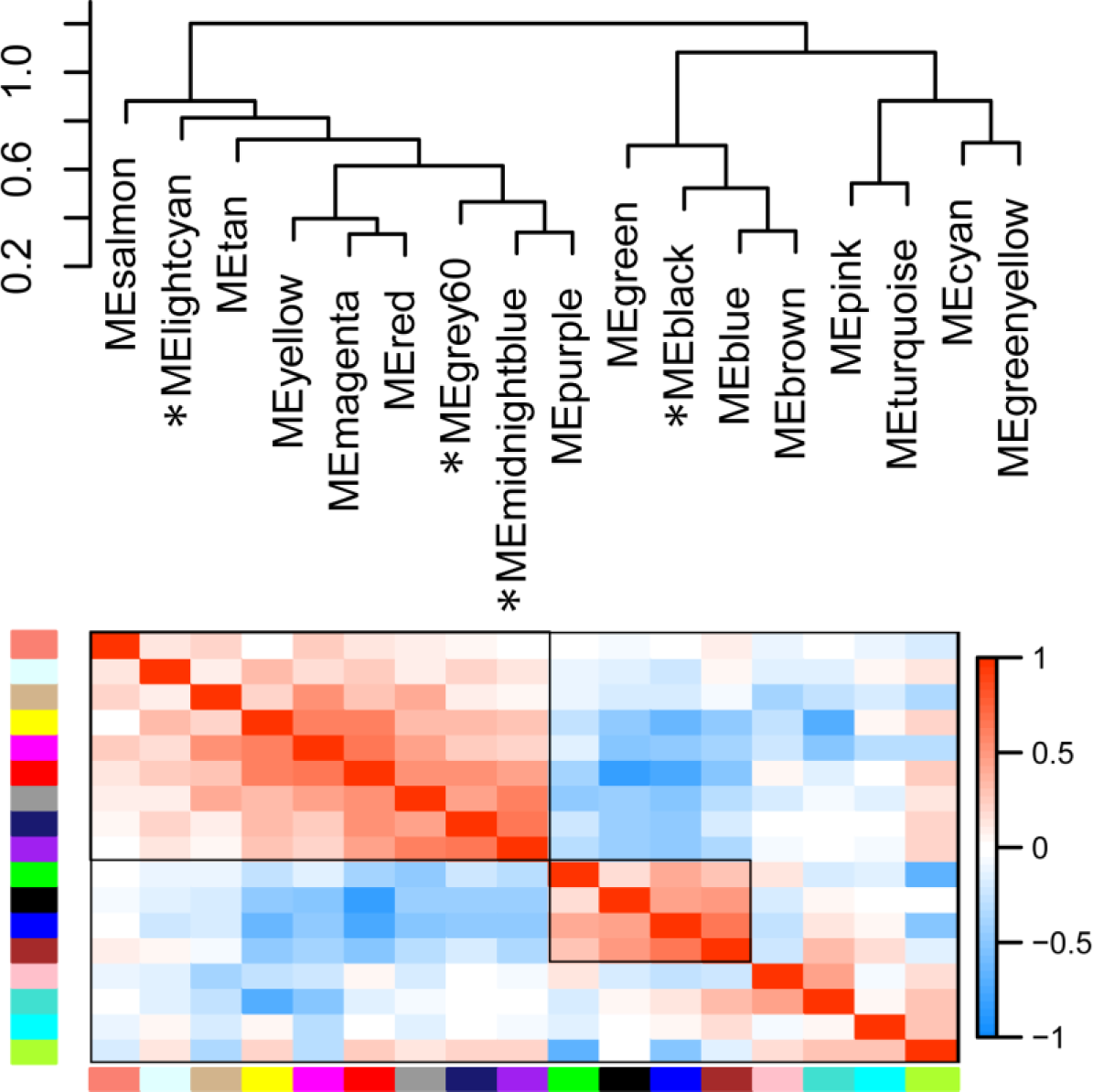
Hierarchical relationships of 17 modules identified in CAMP. Top panel shows eigengene dissimilarity clustering. Bottom panel shows pair-wise Pearson correlations among the eigengenes, ranging from −1 (blue) to 1 (red). The order of the colored columns corresponds to the color names in the top panel. Black boxes denote those modules belonging to the larger and smaller meta-modules. Asterisks denote modules associated with obese asthma in multivariate regression modeling (Table 2). ME, module eigengene.

With 17 modules defined, we next identified ones related to obese asthma by testing for differences in each module’s eigengene distribution between obese and non-obese asthmatics. To obtain a parsimonious differential co-expression model, we performed a systematic parameter selection procedure to prune the multivariate logistic regression model to the most informative subset of corresponding eigengenes, rather than exhaustively testing all possible models [18]. The most parsimonious model (with the lowest BIC) consisted of four eigengenes: black, midnightblue, lightcyan, and grey60 (**Figure E3A**). Three of these eigengenes (midnightblue, lightcyan, and grey60) were part of the larger meta-module, while the fourth (black) was a member of the smaller meta-module (**Figure 2**). Importantly, we observed consistency of module inclusion across all models considered: two of the four eigengenes (black, midnightblue) were represented in greater than 80% of the iterated candidate models, and the other two eigengenes (lightcyan, grey60) were present in greater than 70% of iterated models (**Figure E3B**). Two other models that were within two BIC units of the most parsimonious one each consisted of three of the four modules from that same model: both contained the black and midnightblue eigengenes, but differed in their third members being either lightcyan or grey60. We confirmed that the eigengenes of these four modules (black, midnightblue, lightcyan, grey60) were each strongly and independently associated with adult obesity in the parsimonious model (all *P* < 0.01) (**Table 2**). In contrast to the positive associations of the black, midnightblue, and grey60 eigengenes with adult obesity, the lightcyan eigengene had a negative association. Thus, this model selection procedure converged on a model consisting of four-gene co-expression modules robustly associated with obese asthma.

**Table 2:** Epidemiological eigengene-phenotype model of obese asthma in CAMP. Features are sorted by significance in the adjusted eigengene model. Nominally significant features (*P* < 0.05) are marked in bold. Race and sex were treated as factors, with reference levels of “European” and “Female”, respectively. The race variable from Table 1 was also collapsed into “European” and “Non-European” because of small group sizes. A subset of 112 subjects was missing measured blood count data, but there was no significant difference in expression of any of the four eigengenes between subjects with and without those data (all *P* > 0.1). ME, module eigengene.

| Parameter | Unadjusted Eigengene Model |  |  | Adjusted Eigengene Model |  |  |
| --- | --- | --- | --- | --- | --- | --- |
|  | Beta | SE | P | Beta | SE | P |
| *Modules* |  |  |  |  |  |  |
| MEblack | 12.65 | 3.36 | **1.67E-04** | 15.21 | 4.07 | **1.86E-04** |
| MEmidnightblue | 9.17 | 2.98 | **2.10E-03** | 8.98 | 3.62 | **1.31E-02** |
| MEgrey60 | 7.83 | 2.97 | **8.32E-03** | 8.34 | 3.71 | **2.47E-02** |
| MElightcyan | -7.58 | 2.85 | **7.82E-03** | -4.27 | 6.28 | 4.97E-01 |
| *Covariates (adjusted model only)* |  |  |  |  |  |  |
| Childhood BMI percentile |  |  |  | 0.05 | 0.01 | **7.91E-11** |
| Age at expression |  |  |  | 0.09 | 0.07 | 1.67E-01 |
| Sex |  |  |  | 0.24 | 0.30 | 4.26E-01 |
| % Monocytes |  |  |  | 0.09 | 0.14 | 5.01E-01 |
| % Neutrophils |  |  |  | 0.06 | 0.13 | 6.26E-01 |
| % Eosinophils |  |  |  | 0.06 | 0.15 | 6.68E-01 |
| % Lymphocytes |  |  |  | 0.05 | 0.13 | 7.20E-01 |
| Race |  |  |  | 0.03 | 0.32 | 9.16E-01 |

We examined whether there was evidence of confounding by demographic differences between obese and non-obese CAMP subjects that might affect this parsimonious model. We expanded the unadjusted four-eigengene model to include factors that can be associated with blood expression [12, 23]: age, sex, race, childhood BMI, and blood cell counts. Three of the four eigengenes were still robustly associated with adult obesity in this adjusted epidemiological model of obese asthma (all *P* < 0.01) (**Table 2**). The lightcyan eigengene was very strongly correlated with eosinophil counts (*r* = 0.82. *P* = 9.2E-98) and with neutrophil counts (*r* = −0.24. *P* = 1.1E-06), suggesting that its association with obesity was possibly confounded. Notably, despite its very significant positive association with adult obesity (*P* = 7.9E-11), childhood BMI did not substantively affect the relationships of the three eigengenes with adult obesity. Acting as a negative control, the null eigengene comprising non-co-expressed genes showed no evidence of association with obesity (*P* = 0.52). To facilitate visual understanding of the adjusted obese asthma module model, we also plotted the values of each of the four eigengenes stratified by obesity status (**Figure 3**). Thus, our model selection procedure identified a robust core of gene co-expression modules associated with obese asthma.

**Figure 3:**
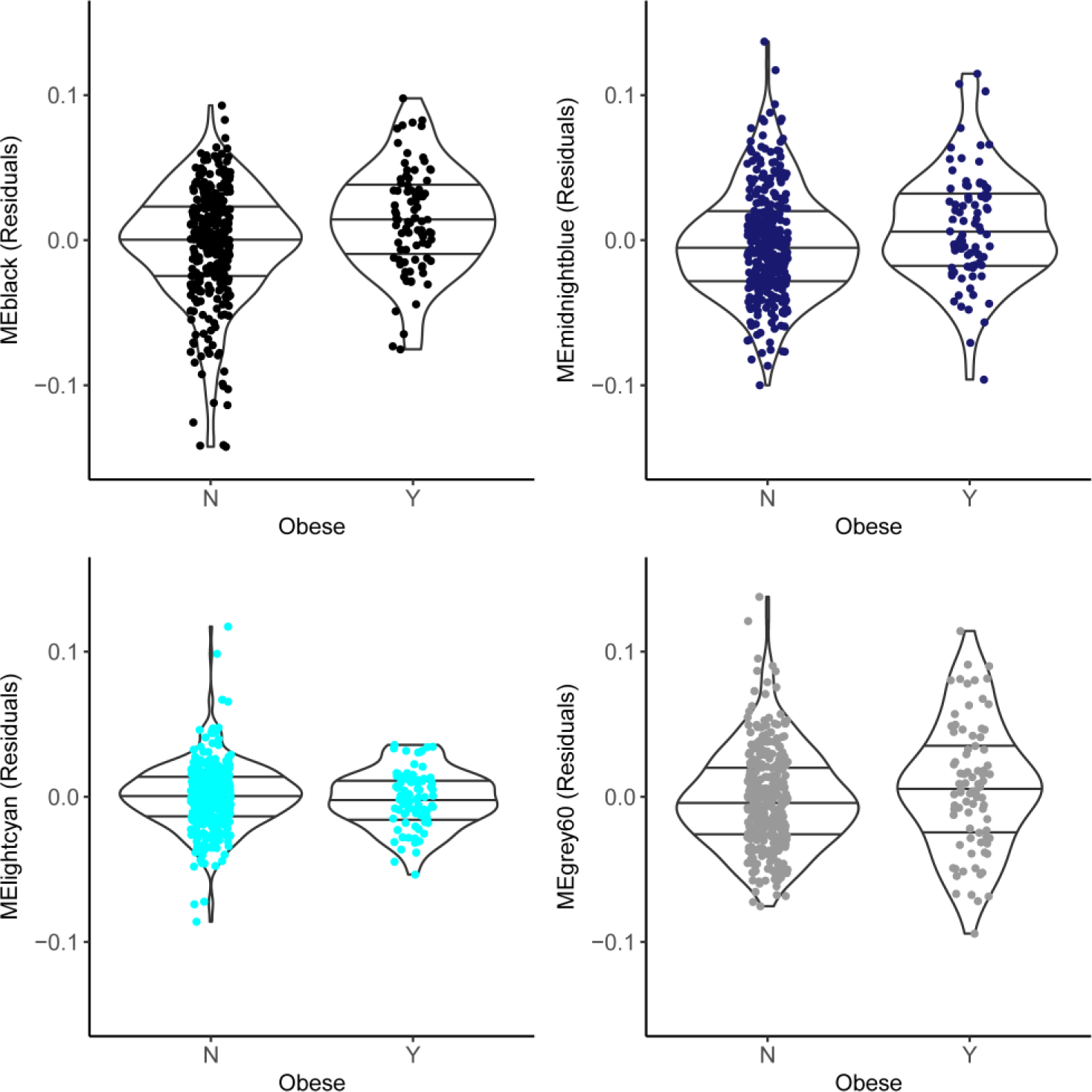
Associations of four eigengenes with obese asthma in CAMP. Panels show violin plots of eigengene residuals by obesity status. Residuals include adjustments for demographic characteristics and the other three eigengenes (see Table 2). Each dot represents an individual CAMP subject. Horizontal lines indicate 25^th^, 50^th^, and 75^th^ percentiles. In this multivariate context, the black, midnightblue, and grey60 eigengenes are significantly different between obese and non-obese asthmatic subjects (*P* < 0.05), while the lightcyan eigengene is not.

### Biological pathways in network modules of obese asthma

To assign biological meaning to interpretability the four obese asthma modules, we performed pathway enrichment analyses on their corresponding gene memberships. We found that the memberships of only two of these modules (black, midnightblue) were significantly enriched (adjusted *P* < 0.05) for genes from biological pathways annotated in the Reactome database (**Table 3**). None of the 48 enriched pathways were shared between these two modules. The grey60 and lightcyan modules had no significantly enriched pathways. Negative control gene sets consisting of 50 or 200 random genes from the null module also had no significantly enriched pathways.

The midnightblue module was enriched for four pathways related to platelet and extracellular matrix biology, and a fifth related to smooth muscle contraction. A subset of 20 of the midnightblue module’s 66 genes (30%) were responsible for the enrichments seen, with six of these genes shared the across at least two pathways. Three of these genes code integrin subunits (*ITGA2B*, *ITGB3*, *ITGB5*), while the others code a cytoskeletal protein (*VCL*), an extracellular matrix protein (*SPARC*), and the Von Willebrand Factor platelet glycoprotein (*VWF*).

The black module was enriched for 44 pathways, including multiple pathways related to mitochondrial and ribosomal biology. These enrichments were driven by 130 of 464 module genes (28%), 24 of which were shared across five or more pathways. A subset of 34 of the enriched pathways involved the same 13 proteasome subunits, including four sets related to non-canonical NF-κB signaling and NF-κB activation (two of which included *NFKB1* itself). Other enriched pathways of interest include those related to interleukin-1 signaling, cystic fibrosis transmembrane conductance regulator (CFTR) dysfunction in cystic fibrosis, and metabolism of the Gli family of transcription factors and other components that all regulate Hh pathway activity.

**Table 3:** Significantly enriched Reactome pathways in two of four eigengenes associated with obese asthma. Term refers to the membership of a given known pathway and query refers to the membership of a given module. Recall is defined as the fraction of genes in the functional term set overlapping the genes in the query set. Precision is defined as the fraction of genes in the query set overlapping the genes in the functional term set.

| Module | Adjusted<br>P Value | Term<br>Size | Query<br>Size | Overlap<br>Size | Precision | Recall | Term ID | Term Name |
| --- | --- | --- | --- | --- | --- | --- | --- | --- |
| midnightblue | 6.32E-08 | 182 | 64 | 16 | 0.25 | 0.088 | REAC:76002 | Platelet activation, signaling and aggregation |
| midnightblue | 0.00119 | 21 | 64 | 5 | 0.078 | 0.238 | REAC:445355 | Smooth muscle contraction |
| midnightblue | 0.0163 | 18 | 64 | 4 | 0.062 | 0.222 | REAC:3000178 | ECM proteoglycans |
| midnightblue | 0.0423 | 9 | 64 | 3 | 0.047 | 0.333 | REAC:372708 | p130Cas linkage to MAPK signaling for integrins |
| black | 6.52E-13 | 76 | 393 | 30 | 0.076 | 0.395 | REAC:5368286 | Mitochondrial translation initiation |
| black | 5.85E-10 | 83 | 393 | 28 | 0.071 | 0.337 | REAC:163200 | Respiratory electron transport, ATP synthesis by chemiosmotic coupling, and heat production by uncoupling proteins. |
| black | 1.20E-04 | 57 | 393 | 17 | 0.043 | 0.298 | REAC:1799339 | SRP-dependent cotranslational protein targeting to membrane |
| black | 1.26E-04 | 70 | 393 | 19 | 0.048 | 0.271 | REAC:450531 | Regulation of mRNA stability by proteins that bind AU-rich elements |
| black | 2.95E-04 | 42 | 393 | 14 | 0.036 | 0.333 | REAC:5362768 | Hh mutants that don't undergo autocatalytic processing are degraded by ERAD |
| black | 3.89E-04 | 119 | 393 | 25 | 0.064 | 0.210 | REAC:72312 | rRNA processing |
| black | 5.24E-04 | 38 | 393 | 13 | 0.033 | 0.342 | REAC:1236978 | Cross-presentation of soluble exogenous antigens (endosomes) |
| black | 5.60E-04 | 44 | 393 | 14 | 0.036 | 0.318 | REAC:8852276 | The role of GTSE1 in G2/M progression after G2 checkpoint |
| black | 7.35E-04 | 39 | 393 | 13 | 0.033 | 0.333 | REAC:69017 | CDK-mediated phosphorylation and removal of Cdc6 |
| black | 7.35E-04 | 39 | 393 | 13 | 0.033 | 0.333 | REAC:69229 | Ubiquitin-dependent degradation of Cyclin D1 |
| black | 7.35E-04 | 39 | 393 | 13 | 0.033 | 0.333 | REAC:211733 | Regulation of activated PAK-2p34 by proteasome mediated degradation |
| black | 7.59E-04 | 45 | 393 | 14 | 0.036 | 0.311 | REAC:187577 | SCF(Skp2)-mediated degradation of p27/p21 |
| black | 7.59E-04 | 45 | 393 | 14 | 0.036 | 0.311 | REAC:5358346 | Hedgehog ligand biogenesis |
| black | 0.00102 | 46 | 393 | 14 | 0.036 | 0.304 | REAC:5678895 | Defective CFTR causes cystic fibrosis |
| black | 0.00102 | 40 | 393 | 13 | 0.033 | 0.325 | REAC:350562 | Regulation of ornithine decarboxylase (ODC) |
| black | 0.00139 | 41 | 393 | 13 | 0.033 | 0.317 | REAC:180534 | Vpu mediated degradation of CD4 |
| black | 0.00139 | 41 | 393 | 13 | 0.033 | 0.317 | REAC:349425 | Autodegradation of the E3 ubiquitin ligase COP1 |
| black | 0.00188 | 42 | 393 | 13 | 0.033 | 0.310 | REAC:4641257 | Degradation of AXIN |
| black | 0.00188 | 42 | 393 | 13 | 0.033 | 0.310 | REAC:174113 | SCF-beta-TrCP mediated degradation of Emi1 |
| black | 0.00252 | 43 | 393 | 13 | 0.033 | 0.302 | REAC:4641258 | Degradation of DVL |
| black | 0.00252 | 43 | 393 | 13 | 0.033 | 0.302 | REAC:180585 | Vif-mediated degradation of APOBEC3G |
| black | 0.00334 | 44 | 393 | 13 | 0.033 | 0.295 | REAC:5610780 | Degradation of GLI1 by the proteasome |
| black | 0.00439 | 45 | 393 | 13 | 0.033 | 0.289 | REAC:5610783 | Degradation of GLI2 by the proteasome |
| black | 0.00439 | 45 | 393 | 13 | 0.033 | 0.289 | REAC:5610785 | GLI3 is processed to GLI3R by the proteasome |
| black | 0.00572 | 46 | 393 | 13 | 0.033 | 0.283 | REAC:4608870 | Asymmetric localization of PCP proteins |
| black | 0.00722 | 183 | 393 | 30 | 0.076 | 0.164 | REAC:71291 | Metabolism of amino acids and derivatives |
| black | 0.00739 | 47 | 393 | 13 | 0.033 | 0.277 | REAC:5676590 | NIK-->noncanonical NF-kB signaling |
| black | 0.0080 | 54 | 393 | 14 | 0.036 | 0.259 | REAC:1169091 | Activation of NF-kappaB in B cells |
| black | 0.00881 | 260 | 393 | 38 | 0.097 | 0.146 | REAC:5663205 | Infectious disease |
| black | 0.00947 | 48 | 393 | 13 | 0.033 | 0.271 | REAC:5607761 | Dectin-1 mediated noncanonical NF-kB signaling |
| black | 0.00947 | 48 | 393 | 13 | 0.033 | 0.271 | REAC:1234176 | Oxygen-dependent proline hydroxylation of<br>Hypoxia-inducible Factor Alpha |
| black | 0.012 | 49 | 393 | 13 | 0.033 | 0.265 | REAC:72689 | Formation of a pool of free 40S subunits |
| black | 0.012 | 49 | 393 | 13 | 0.033 | 0.265 | REAC:5658442 | Regulation of RAS by GAPs |
| black | 0.012 | 49 | 393 | 13 | 0.033 | 0.265 | REAC:174084 | Autodegradation of Cdh1 by Cdh1:APC/C |
| black | 0.014 | 71 | 393 | 16 | 0.041 | 0.225 | REAC:1236974 | ER-Phagosome pathway |
| black | 0.0154 | 37 | 393 | 11 | 0.028 | 0.297 | REAC:192823 | Viral mRNA Translation |
| black | 0.0238 | 52 | 393 | 13 | 0.033 | 0.250 | REAC:179419 | APC/Cdc20 mediated degradation of cell cycle<br>proteins prior to satisfaction of the cell cycle<br>checkpoint |
| black | 0.0264 | 39 | 393 | 11 | 0.028 | 0.282 | REAC:2408557 | Selenocysteine synthesis |
| black | 0.0285 | 75 | 393 | 16 | 0.041 | 0.213 | REAC:446652 | Interleukin-1 signaling |
| black | 0.0294 | 53 | 393 | 13 | 0.033 | 0.245 | REAC:5632684 | Hedgehog 'on' state |
| black | 0.0362 | 54 | 393 | 13 | 0.033 | 0.241 | REAC:156827 | L13a-mediated translational silencing of<br>Ceruloplasmin expression |
| black | 0.0416 | 62 | 393 | 14 | 0.036 | 0.226 | REAC:2871837 | FCERI mediated NF-kB activation |
| black | 0.0436 | 41 | 393 | 11 | 0.028 | 0.268 | REAC:975956 | Nonsense Mediated Decay (NMD) independent of<br>the Exon Junction Complex (EJC) |
| black | 0.0444 | 55 | 393 | 13 | 0.033 | 0.236 | REAC:174178 | APC/C:Cdh1 mediated degradation of Cdc20 and<br>other APC/C:Cdh1 targeted proteins in late<br>mitosis/early G1 |

### Preservation and replication of obese asthma modules

To assess the reproducibility and generalizability of our results, we replicated our analysis in two independent datasets. One replication cohort consisted of 107 asthmatic adults (CHS) (**Table E3**) and the other consisted of 327 asthmatic children (GACRS) (**Table E4**). Obese and non-obese subjects in CHS differed from each other in terms of childhood BMI (*P* < 0.05), but not in terms of age of asthma onset, age at gene expression, proportion of females, ethnicity, or blood cell counts (*P* > 0.1). Obese and non-obese subjects in GACRS did not significantly differ from each other in terms of age of asthma onset, age at gene expression, proportion of females, ethnicity, IgE level, or blood cell counts (*P* > 0.1).

Replication analysis in CHS and GACRS consisted of two steps: module preservation and module association. Preservation refers to the consistency of network module structure across datasets. If modules are not preserved across cohorts, then their corresponding eigengenes are not comparable, thus precluding further eigengene-phenotype association testing. The seven preservation statistics are motivated in detail elsewhere [8, 21, 24].

We found very strong evidence of module preservation in the two replication cohorts. All four modules from CAMP were highly significantly preserved in both CHS and GARCS (*P* = 9.999E-05 for each, based on permutation testing, **Tables E5** and **E6**). The seven preservation statistics for each of the modules were very similar in magnitude between and within the two replication cohorts. Visualization of the four modules’ properties in the replication cohorts further reinforced the notion of strong preservation (**Figure 4**). Within each module, we observed four patterns. First, genes were consistently positively correlated with each other but not with genes in other modules. Second, the vast majority of gene-pairs had edge weights close to zero, reflecting that only a minority of genes were strongly correlated with more than a few other genes. Third, those same genes with higher edge weights had higher weighted degrees, reflecting their increased network connectivity. In other words, genes that were highly connected (“hub-like”) in the discovery cohort also acted as module hubs in the replication cohorts. Fourth, node contributions were broadly directionally consistent, meaning that genes were consistently correlated with the corresponding module summary measure (i.e., the eigengene). Thus, the obese asthma network modules identified in CAMP had reproducible architectures in CHS and GACRS.

**Figure 4:**
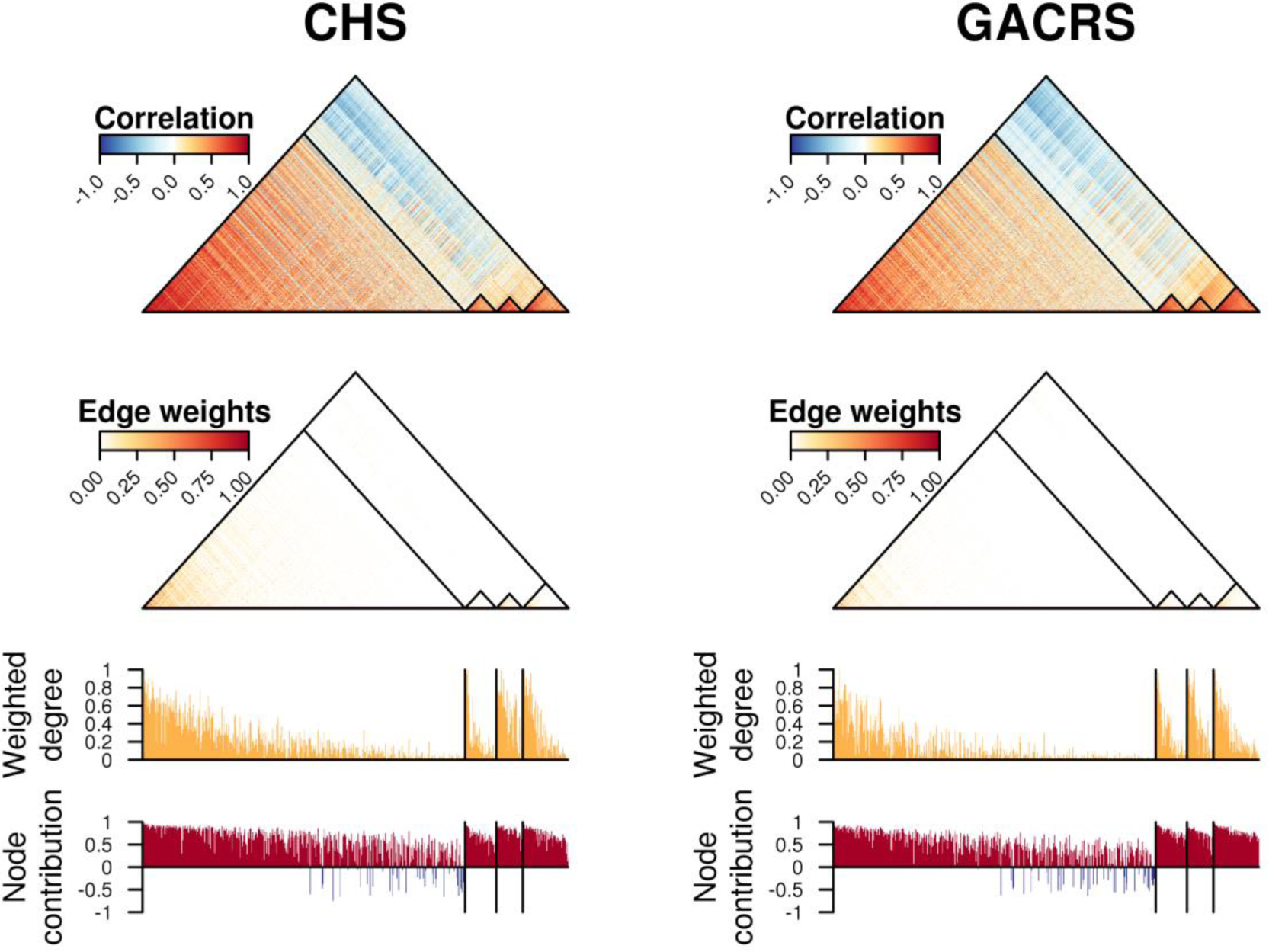
Preservation of obese asthma modules in CHS and GACRS. The first (top) panel shows a heatmap of pair-wise correlations among the genes comprising the black, midnightblue, grey60, and lightcyan modules [see ‘cor.cor’ and ‘avg.cor’ metrics in Table E5]. The second panel shows a heatmap of the edge weights (connections) among the genes comprising the four modules [‘avg.weight’]. The third panel shows the distribution of scaled weight degrees (relative connectedness) among the genes comprising the four modules [‘cor.degree’]. The fourth panel shows the distribution of node contributions (correlation to module eigengene) among the genes comprising the four modules [‘cor.contrib’ and ‘avg.contrib’]. The ‘coherence’ metric is not summarized in this figure. Genes are ordered from left to right based on their weighted degree in CAMP (the discovery cohort) to highlight the consistency of the network properties in CHS and GACRS (the replication cohorts).

In the second stage of replication, we tested these highly preserved modules for evidence of phenotypic association with obesity in the two additional asthma cohorts. Testing was done at two levels: (i) assessing replication of each module’s eigengene association with obese asthma; and (ii) assessing replication at the gene level by GSEA of differential expression of each module’s individual component genes. Minimal evidence for replication was observed at the eigengene level: no significant associations were observed in CHS, and although in GARCS the midnightblue eigengene was significantly associated with BMI *Z*-score, its direction of effect was opposite of that observed in the CAMP discovery cohort (**Tables E7** and **E8**).

In contrast, we found stronger replication evidence when considering individual expression differences for each module’s gene components using GSEA. In CAMP, all module gene sets except those in the lightcyan module were significantly enriched for differential expression between obesity and non-obese asthmatics (*P* < 0.05) (**Table 4**), in directions consistent with their modules’ original eigengene associations (**Table 2**). In CHS, the enrichments for three of the four modules were similarly significant in the direction consistent with the CAMP results. In GACRS, the enrichments for all four modules were significant in the direction consistent with the CAMP results. Furthermore, negative control gene sets consisting of 50 or 200 random genes from the null module were generally not significantly enriched in any of the three cohorts (*P* > 0.2). These gene set enrichment results paralleled the fact that in each of the four modules, the node contributions of the corresponding genes were very consistent across the three cohorts (**Figure 4**). These complementary differential gene expression results further supported the existence of four co-expression modules associated with obese asthma.

**Table 4:**
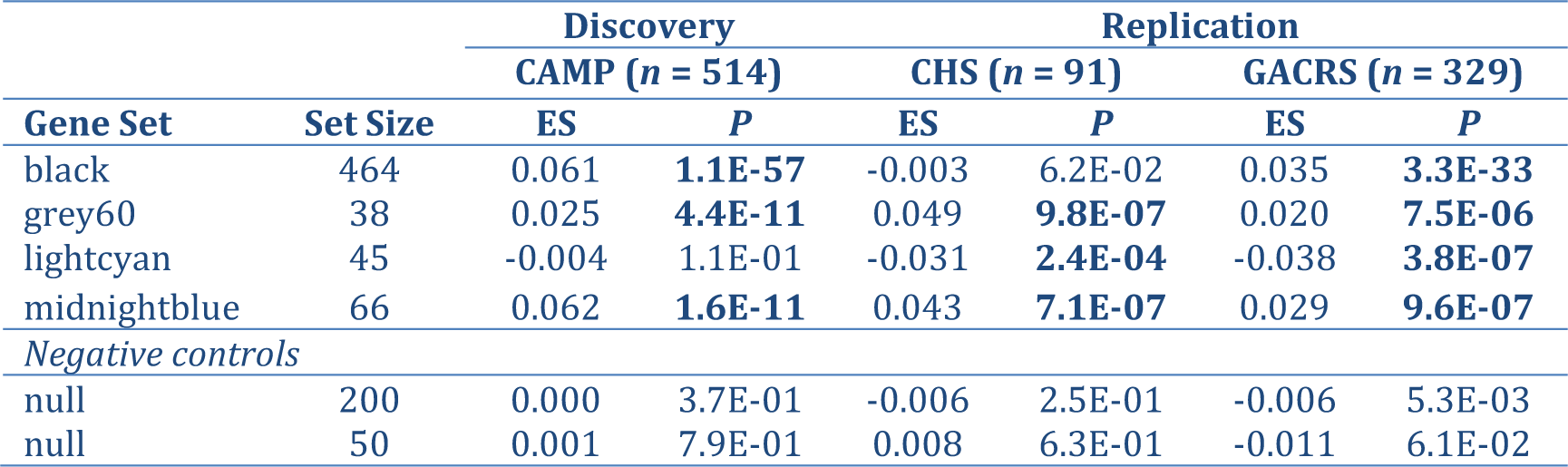
Enrichments for differential expression of obese asthma module gene sets. Significant enrichments (*P* < 0.05) with a consistent direction of effect to results from Table 2 are marked in bold. ES, enrichment score. Two random subsets of the null module were used as negative control sets.

## Discussion

In this work, we used whole blood gene expression profiling to characterize the biology of a form of obese asthma known as “asthma complicated by obesity” [2] (i.e., cases where asthma preceded the development of obesity). Among 514 subjects who were diagnosed with asthma as children, we identified four robust co-expression network modules jointly associated with adult obesity. We then demonstrated compelling statistical evidence of replication in a total of 418 subjects from two independent asthma cohorts (one adult and one pediatric). Two of these obese asthma networks, termed the “black” and “midnightblue” modules, were significantly enriched for genes from several inflammatory biological pathways of interest (e.g., platelets, smooth muscle contraction, integrins, NF-κB signaling).

Pre-existing scientific evidence strongly supports the disease relevance of our obese asthma expression signatures. The most strongly enriched pathway for the midnightblue co-expression module comprised genes related to “platelet activation, signaling and aggregation”. Several studies have linked platelets with allergic airway disease through both innate and adaptive immune pathways affecting bronchoconstriction, airway remodeling, and airway inflammation [25, 26]. In mice, the lungs have been shown to be a major site of platelet biogenesis [27]. More apt to our findings, several studies have demonstrated important links between platelets and obese asthma. In a published electronic medical record review, asthmatic children who were obese had higher platelet counts than those who were lean [28]; a comparable relationship between BMI and platelet counts was observed in adults [29]. Weight loss due to bariatric surgery was shown to reduce mean platelet volume concurrently with other inflammatory biomarkers [30]. These data, together with our results, strongly support a role of platelet biology in obese asthma that merits further investigation.

The midnightblue module was also enriched for pathways related to integrins and smooth muscle contraction. Integrins are transmembrane proteins that anchor airway smooth muscle to its extracellular matrix that may be important novel therapeutic targets in asthma because of their roles in airway hyper-responsiveness and remodeling [31]. That a gene module associated with obese asthma is enriched for these two molecular pathways may be especially important, potentially identifying molecular processes that contribute to impaired lung function [32], increased airway obstruction [10], and reduced responsiveness to corticosteroid [33] and bronchodilator therapies [34] observed in obese asthmatics. Supporting this notion, *in vitro* suppression of eosinophil-derived integrins prevented airway smooth muscle remodeling in samples from asthmatic individuals (of unknown BMI) [35]. Airway smooth muscle function has also been shown to be affected by consumption of a high-fat diet [36], a key exposure in the development of obesity. Finally, integrins mediate the interaction of activated platelets and eosinophils in relation to asthma severity [37].

The black module was dominated by a ribosome and proteasome-related gene signature. Interestingly, similar signatures have been reported in two blood expression studies of obesity unrelated to asthma: a ribosome pathway signature defined in peripheral whole blood samples was also associated with obesity in a small gene expression study in 17 obese and 17 non-obese subjects [38], and a second study by the same group showed a proteasome signature in whole blood comparing nine diet-sensitive and nine diet-resistant obese subjects [39]. Given that both these biological processes are of fundamental importance to cellular function generally, it is unclear whether their implication in our study is specific to asthma and obesity together (perhaps reflecting fundamental differences in cellular metabolic demands in these patients) [40, 41], or whether these associations reflect more generic patterns of cellular activity or stress seen in obese patients [42, 43].

The black module was also enriched for a quartet of very closely related NF-κB signaling and activation pathways, which are highly relevant to the biology of obese asthma. A mouse model of obese asthma has suggested that the pathobiology includes increased NF-κB signaling and oxidative stress in lung tissue [44]. Furthermore, metformin has been shown to inhibit the TNF-α-induced inflammatory signaling and NF-κB-mediated iNOS expression in lung tissue of obese mice [45], suggesting the drug could attenuate the exacerbation of allergic eosinophilic inflammation in obese asthma. In humans, metformin has also been shown to reduce risk of asthma-related outcomes among individuals with both asthma and diabetes [46]. Thus, NF-κB signaling is an intriguing candidate, including potential therapeutic consideration, for further functional study in the context of obese asthma and potential therapeutic targeting.

Finally, the black module was also enriched for pathways related to factors regulating Hh pathway activity, such as interleukin-1 (IL-1), CFTR, and the Gli family of transcription factors. Broadly, Hh signaling is a developmental process linked to both adipogenesis [47] and airway remodeling [48], and its regulatory factors are linked to asthma. Airway gene expression profiling in adults with obstructive airway diseases demonstrated the associations of a set of IL-1 pathway mediators with exacerbation risk [49]. CFTR is also expressed in airway smooth muscle cells and has a putative functional role in bronchodilation [50]. Gli and Sonic Hh signaling have been shown to enhance the Th2 immune response in a murine model of allergic asthma [51]. It is known that Hh signaling serves a similar function to Wnt signaling [48, 52], particularly in relation to airway remodeling. Fetal lung gene expression profiling has implicated Wnt receptor signaling in the developmental programming of airway branching morphogenesis and of downstream childhood asthma risk [53]. Such mechanisms affecting lung structure are also highly relevant to our understanding of obese asthma [2].

Despite the overall robustness of our study findings, there are important epidemiological, genomic, and statistical limitations to consider. It is possible that our findings may not be applicable to other endotypes of obese asthma, such as when the development of obesity precedes the development of asthma rather than the other way around, as in the cohorts we studied. That our findings replicate in two cross-sectional cohorts, where the order of asthma and obesity occurrence is not known, suggests that our signatures may indeed apply to other obese asthma endotypes; corroboration of this in additional cohorts is needed. Similarly, because our study did not include contrasts with non-asthmatic controls with and without obesity, we cannot definitively conclude whether the observed signatures are specific to obese asthma, or would also be seen in obese individuals without asthma. We also note the lack of significant pathway enrichments for the “lightcyan” and “grey60” modules. One possible explanation is that the underlying biology represented by those two modules was not well characterized as coherent pathways in Reactome.

We also recognize that both asthma and obesity are multi-system disorders, involving the coordinated interactions of a variety of cellular compartments. Our focus on patterns of gene expression in whole blood samples is unlikely to capture molecular processes that are specific to other relevant tissues. For this, ideally, we would need to develop co-expression networks across multiple disease-relevant tissues (adipose, lung, and blood) in the same set of individuals. One such example in the literature is of cross-sectional expression data in liver, omental adipose, subcutaneous adipose, and stomach obtained from morbidly obese individuals undergoing bariatric surgery [54]. Unfortunately, such data in a relatively large number of subjects do not currently exist for the study of obese asthma. Furthermore, peripheral blood by itself is a very heterogeneous tissue, though we used differential cell count measurements to account for this effect on expression profiles in our association models. Despite these limitations, measuring gene expression only in blood does provide an accessible initial route to insights into the biology of obese asthma, particularly given the importance of the peripheral blood in facilitating communication between adipose tissue, the lung, and immune compartments.

Finally, WGCNA produces gene co-expression networks that characterize the undirected correlational relationships between genes. Because WGCNA-derived networks are undirected, they are therefore not informative about directed causal relationships (i.e., which gene in a correlated pair is upstream of the other in a biological pathway). Though this could be addressed somewhat through the analysis of longitudinal gene expression data (measured before and after the development of obesity), such information was not available for this study.

## Conclusion

In summary, we have identified four robust gene co-expression network modules of asthma complicated by obesity. Nearly every module-related biological pathway we highlighted has been implicated in both asthma and obesity. Together, these findings suggest a few mechanistic possibilities: (1) that subsets of individuals might be genetically programmed with shared pathways that lead to this obese asthma phenotype; or (2) that a subset of asthmatic individuals has a second set of exposures that begets changes in expression resulting in consequent obesity; or (3)that the subsequent development of obesity itself triggers the observed expression change. We propose these candidate genes and pathways merit further examination in ongoing and future epidemiological studies and clinical trials of obese asthma.

## Acknowledgements

We thank the Asthma BRIDGE Consortium study staff for their assistance and contributions, as well as all the study subjects for their participation in the Asthma BRIDGE initiative.

## Author Contributions

- D.C.C.-C. and B.A.R. designed the study.
- D.C.C.-C. performed the analyses.
- K.C.B., A.B.-V., J.C.C., W.J.G., F.D.G., J.A.K., A.H.L., S.J.L., F.D.M., J.M., E.T.N., D.L.N., S.R.W., C.O., S.T.W., and B.A.R. supervised the primary data generation.
- Z.C. curated additional phenotype data for analysis.
- D.C.C.-C. and B.A.R. wrote the manuscript.
- All co-authors read and approved the final manuscript.

## Supplementary Material

**Figure E1:**
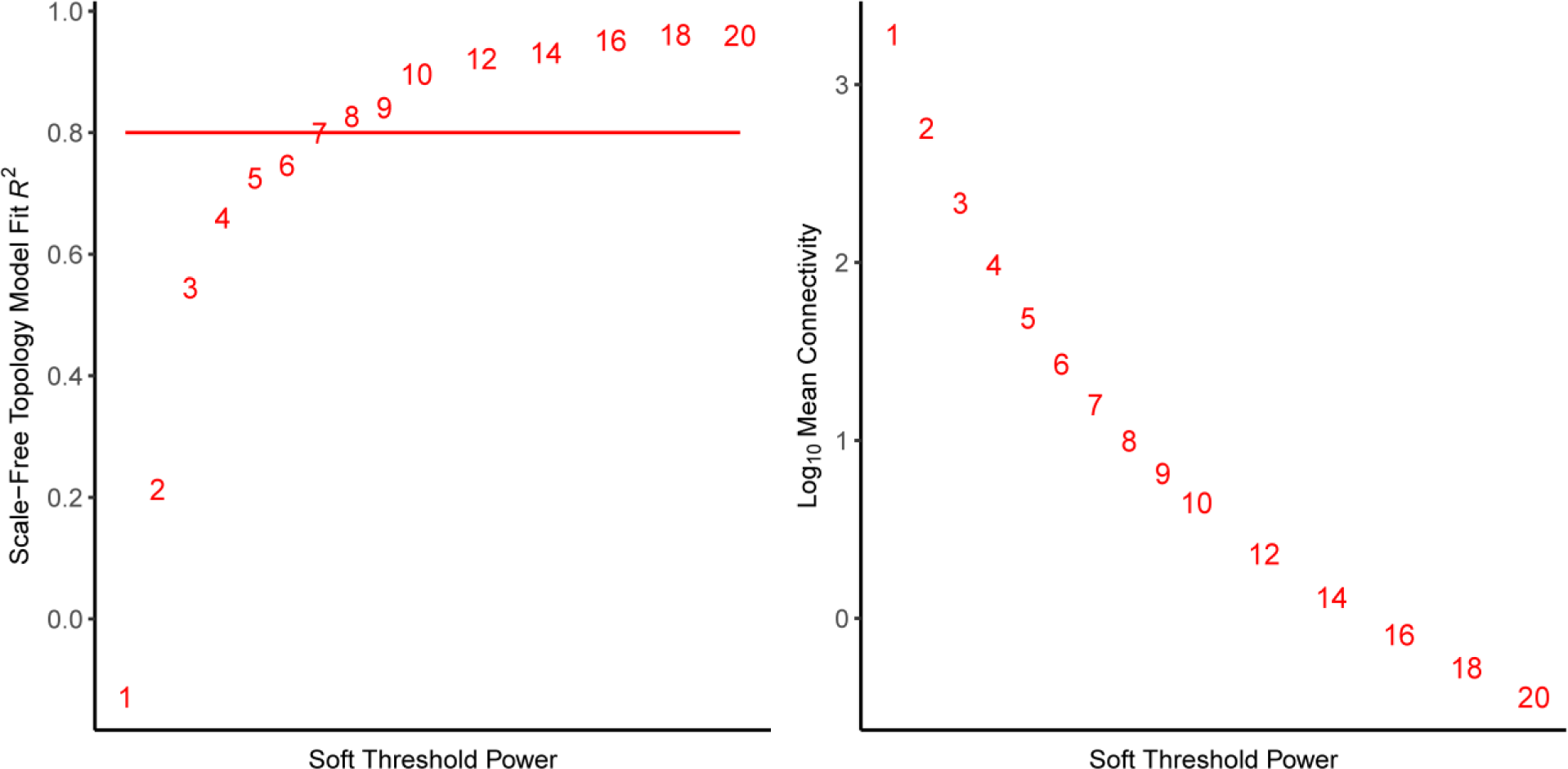
Assessing scale-free model fitting in CAMP. Left panel shows scale-free topology plotted by soft threshold. The red horizontal line represents the cutoff for identifying a strong model fit. Right panel shows mean gene connectivity plotted by soft threshold.

**Figure E2:**
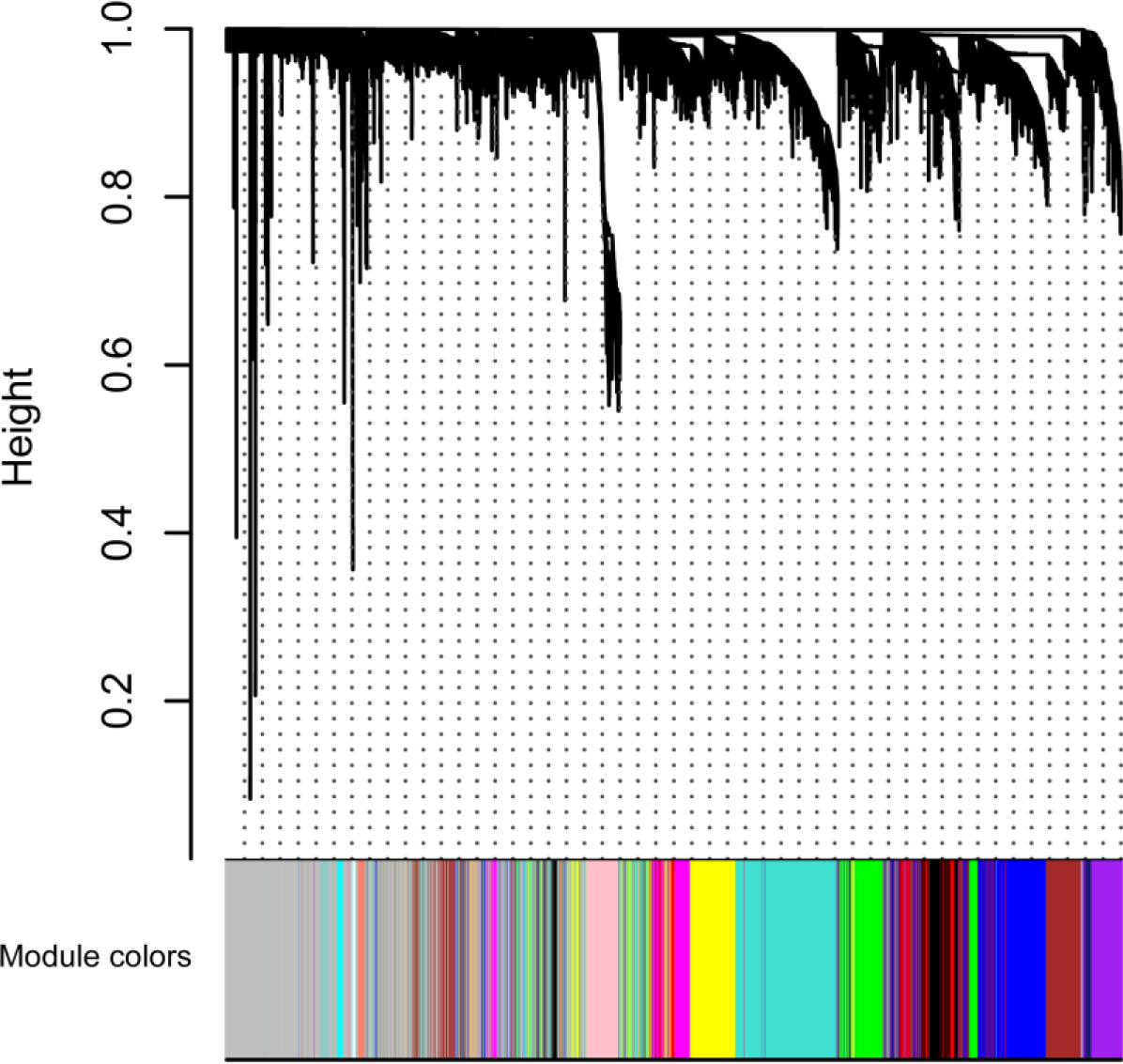
Gene co-expression clustering in CAMP. Top panel shows a Dendrogram of 10,448 genes. Bottom panel shows colors corresponding to the cluster membership labels for each gene (see Tables E1 and E2).

**Table E1:** Sizes of 17 gene co-expression network modules identified in CAMP. Each module was assigned an arbitrary color label. The null module is a collection of 2,670 genes that could not be assigned membership to a module because of insufficient co-expression. Each assayed gene could only be a member of a single module. Module memberships are listed in Table E2. Asterisks denote modules associated with obese asthma in multivariate regression modeling (Table 2). N, number of genes in each module.

| Module Name | N | Module Name | N |
| --- | --- | --- | --- |
| turquoise | 1,686 | purple | 348 |
| blue | 1,091 | greenyellow | 116 |
| brown | 971 | tan | 114 |
| yellow | 825 | salmon | 82 |
| green | 565 | cyan | 69 |
| red | 469 | midnightblue* | 66 |
| black* | 464 | lightcyan* | 45 |
| pink | 431 | grey60* | 38 |
| magenta | 398 |  |  |

**Table E2:**
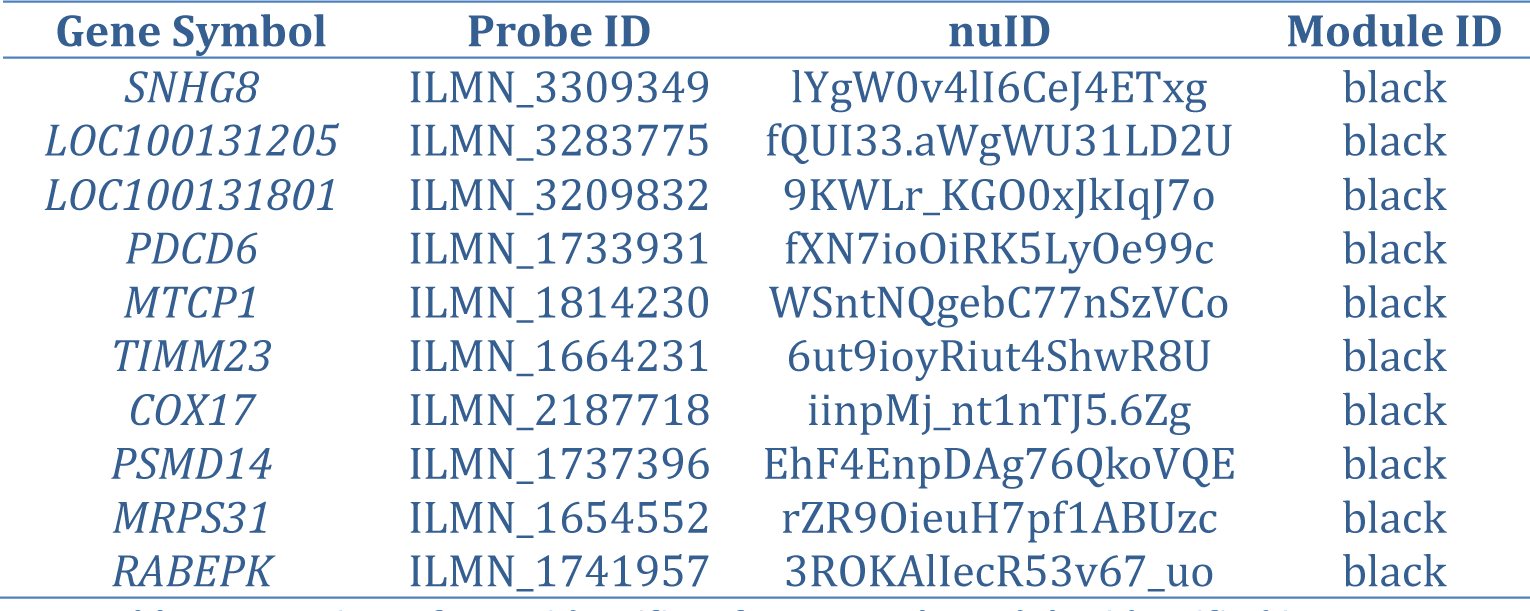
Preview of gene identifiers for network modules identified in CAMP. A complete list of all 10,448 IDs is provided as an external text file. nuID, nucleotide universal identifier.

**Figure E3:**
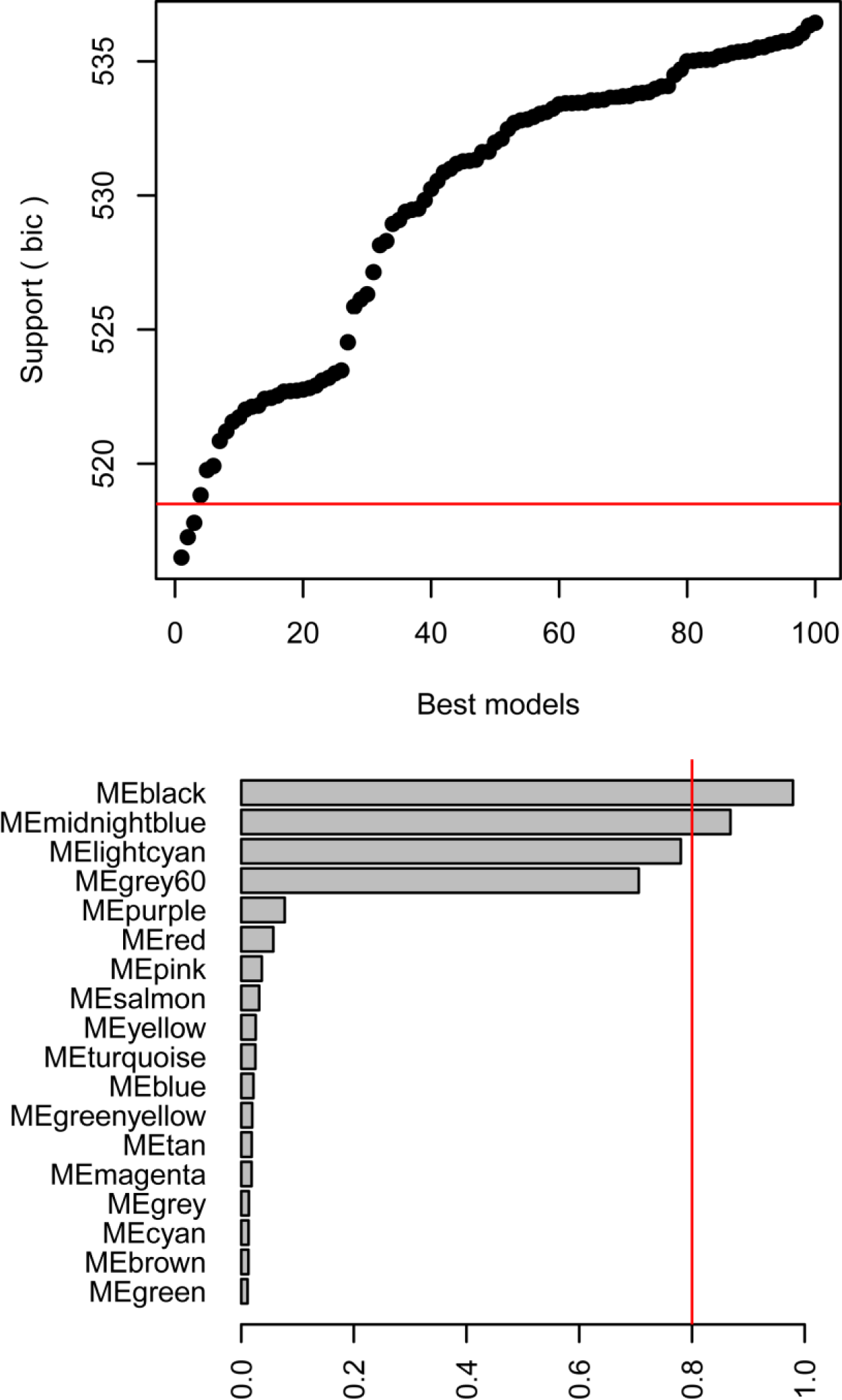
Eigengene-phenotype model selection in CAMP. Top panel (A) shows the distribution of BIC values among the 100 best eigengene models of obese asthma generated by the model selection procedure. A horizontal red line indicates that models below it that are comparable to the best model (less than 2 BIC units away from each other). Bottom panel (B) shows the model-averaged importance of terms across all models generated by the selection procedure. A vertical red line indicates which eigengenes were present in 80% or more of iterated models. ME, module eigengene.

**Table E3:** CHS replication cohort characteristics. To test for differences between obese and non-obese subjects, Welch two sample *t*-tests were used for continuous traits, and chi-squared tests were used for proportions. Blood cell percentages were estimated based on gene expression, except for eosinophils, whose levels were not measured directly and could not be imputed.

|  | **Obese** | **Non-Obese** | **Difference *P*** |
| --- | --- | --- | --- |
| *N* | 18 | 73 | - |
| **Childhood BMI percentile** | 77.6 17.8 | 55.4 28.7 | 1.7E-04 |
| **Adult BMI** | 32.8 (2.6) | 24.6 (3.4) | 6.93E-13 |
| **Age at asthma onset (S.D.) [yrs.]** | 5.6 (4.1) | 7.0 (4.9) | 0.23 |
| **Age at gene expression (S.D.) [yrs.]** | 26.6 (2.2) | 27.2 (2.1) | 0.99 |
| **% Female** | 50.0 | 67.1 | 1.00 |
| **Race/Ethnicity *N* (%)** |  |  |  |
| **European** | 4 (22.2) | 27 (37.0) | 0.088 |
| **Black/African American** | 0 (0.0) | 0 (0.0) |  |
| **Hispanic/Latino** | 11 (61.1) | 24 (32.9) |  |
| **Other** | 3 (16.7) | 22 (30.1) |  |
| **% Eosinophils (S.D.)** | - | - | - |
| **% Lymphocytes (S.D.)** | 33.8 (4.6) | 33.2 (4.2) | 0.66 |
| **% Monocytes (S.D.)** | 9.0 (1.1) | 8.8 (1.3) | 0.48 |
| **% Neutrophils (S.D.)** | 47.8 (6.1) | 47.9 (5.2) | 0.062 |

**Table E4:** GACRS replication cohort characteristics. To test for differences between obese and non-obese subjects, Welch two sample *t*-tests were used for continuous traits, and chi-squared tests were used for proportions. Race is not reported as all subjects are Costa Rican. Childhood obesity was defined as a BMI ≥ 95th percentile. Blood cell percentages were estimated based on gene expression, except for eosinophils, whose levels were measured directly.

|  | **Obese** | **Non-Obese** | **Difference *P*** |
| --- | --- | --- | --- |
| *N* | 53 | 274 | - |
| **Childhood BMI percentile** | 97.4 (1.3) | 52.8 (29.1) | - |
| **Age at asthma onset (S.D.) [yrs.]** | 0.3 (0.5) | 0.3 (0.6) | 0.49 |
| **Age at gene expression (S.D.) [yrs.]** | 9.4 (1.8) | 9.0 (1.7) | 0.13 |
| **% Female** | 39.6 | 43.4 | 1.00 |
| **Log<sub>10</sub> IgE** | 2.5 (0.6) | 2.5 (0.7) | 0.78 |
| **Log<sub>10</sub> eosinophil counts (S.D.)** | 2.5 (0.4) | 2.5 (0.5) | 0.24 |
| **% Lymphocytes (S.D.)** | 40.5 (5.5) | 40.1 (4.9) | 0.65 |
| **% Monocytes (S.D.)** | 6.7 (1.0) | 6.6 (1.0) | 0.50 |
| **% Neutrophils (S.D.)** | 45.7 (5.0) | 45.9 (4.6) | 0.79 |

**Table E5:** Module preservation statistics in CHS. All metrics were highly significant after permutation testing (all maximum *P* = 9.999E-05). Columns correspond to seven module preservation statistics (definitions from NetRep package vignette [21]): ‘cor.cor’ measures the concordance of the correlation structure: or, how similar the correlation heatmaps are between the two datasets. ‘avg.cor’ measures the average magnitude of the correlation coefficients of the module in the test dataset: or, how tightly correlated the module is on average in the test dataset. This score is penalized where the correlation coefficients change in sign between the two datasets. ‘avg.weight’ measures the average magnitude of edge weights in the test dataset: or how connected nodes in the module are to each other on average. ‘cor.degree’ measures the concordance of the weighted degree of nodes between the two datasets: or, whether the nodes that are most strongly connected in the discovery dataset remain the most strongly connected in the test dataset. ‘cor.contrib’ measures the concordance of the node contribution between the two datasets: this measures whether the module’s summary profile summarizes the data in the same way in both datasets. ‘avg.contrib’ measures the average magnitude of the node contribution in the test dataset: this is a measure of how coherent the data is in the test dataset. This score is penalized where the node contribution changes in sign between the two datasets: for example, where a gene is differentially expressed between the two datasets. ‘coherence’ measures the proportion of variance in the module data explained by the module’s summary profile vector in the test dataset.

| Module | avg.weight | coherence | cor.cor | cor.degree | cor.contrib | avg.cor | avg.contrib |
| --- | --- | --- | --- | --- | --- | --- | --- |
| black | 0.02 | 0.42 | 0.78 | 0.88 | 0.86 | 0.36 | 0.60 |
| midnightblue | 0.02 | 0.50 | 0.67 | 0.84 | 0.72 | 0.47 | 0.69 |
| grey60 | 0.07 | 0.64 | 0.68 | 0.71 | 0.63 | 0.62 | 0.79 |
| lightcyan | 0.02 | 0.48 | 0.69 | 0.90 | 0.81 | 0.44 | 0.67 |

**Table E6:**
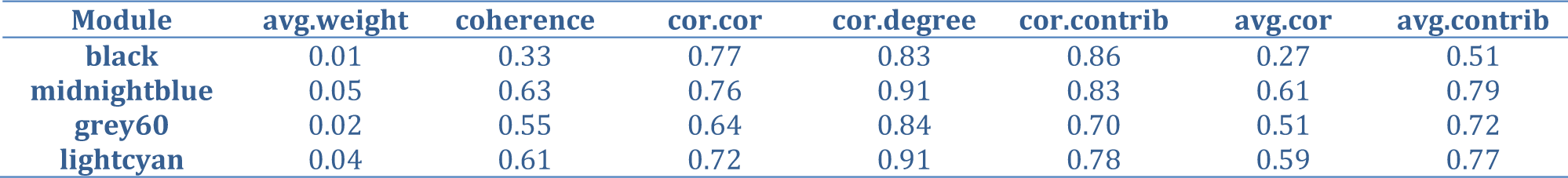
Module preservation statistics in GACRS. All metrics were highly significant after permutation testing (all maximum *P* = 9.999E-05). Columns correspond to seven module preservation statistics defined in Table E5.

**Table E7:** Epidemiological eigengene-phenotype model of obese asthma in CHS. Features are sorted by significance. Nominally significant features (*P* < 0.05) are marked in bold. Race and sex were treated as factors and base levels were “European” and “Female”, respectively. ME, module eigengene.

| Parameter | Beta | SE | P |
| --- | --- | --- | --- |
| **Childhood BMI percentile** | 0.05 | 0.02 | **7.48E-03** |
| % Monocytes | 0.61 | 0.39 | 1.18E-01 |
| % Lymphocytes | 0.21 | 0.17 | 2.24E-01 |
| Sex | 0.83 | 0.70 | 2.36E-01 |
| MEmidnightblue | -8.15 | 7.35 | 2.68E-01 |
| MElightcyan | -10.62 | 9.58 | 2.68E-01 |
| % Neutrophils | 0.15 | 0.15 | 3.03E-01 |
| Race (Hispanic/Latino) | 0.79 | 0.85 | 3.52E-01 |
| MEblack | 7.56 | 15.58 | 6.27E-01 |
| MEgrey60 | 5.52 | 12.73 | 6.65E-01 |
| Race (Other) | 0.13 | 0.93 | 8.86E-01 |
| Age at expression | -0.01 | 0.16 | 9.34E-01 |

**Table E8:** Epidemiological eigengene-phenotype model of obese asthma in GACRS. Features are sorted by significance. Nominally significant features (*P* < 0.05) are marked in bold. Race was treated as a factor and the base level was “Female”. ME, module eigengene.

| Parameter | Beta | SE | P |
| --- | --- | --- | --- |
| **MEmidnightblue** | -8.86 | 4.40 | **4.47E-02** |
| MEblack | 13.50 | 6.87 | 5.04E-02 |
| Age at expression | 0.07 | 0.04 | 6.63E-02 |
| MEgrey60 | -5.24 | 4.33 | 2.27E-01 |
| % Neutrophils | 0.03 | 0.06 | 5.77E-01 |
| % Lymphocytes | 0.03 | 0.06 | 6.29E-01 |
| Sex | -0.04 | 0.14 | 7.87E-01 |
| Log<sub>10</sub> eosinophils | 0.03 | 0.14 | 8.32E-01 |
| MElightcyan | -0.57 | 3.22 | 8.60E-01 |
| % Monocytes | -0.01 | 0.09 | 9.11E-01 |

